# Influence of sequencing depth on the fidelity and sensitivity of 1%-5% low-frequency mutation detection and recommendation for standardization of sequencing depth

**DOI:** 10.1101/2021.12.26.474219

**Authors:** Zhe Liu, Weijin Qiu, Shujin Fu, Xia Zhao, Jun Xia, Chunyu Geng, Youqian Yu, Ziling Li, Mingzhu Li, Hui Jiang, Fang Chen

## Abstract

Sequencing depth has always played an important role in the accurate detection of low-frequency mutations. The increase of sequencing depth and the reasonable setting of threshold can maximize the probability of true positive mutation, or sensitivity. Here, we found that when the threshold was set as a fixed number of positive mutated reads, the probability of both true and false-positive mutations increased with depth. However, When the number of positive mutated reads increased in an equal proportion with depth (the threshold was transformed from a fixed number to a fixed percentage of mutated reads), the true positive probability still increased while false positive probability decreased. Through binomial distribution simulation and experimental test, it is found that the “fidelity” of detected-VAFs is the cause of this phenomenon. Firstly, we used the binomial distribution to construct a model that can easily calculate the relationship between sequencing depth and probability of true positive (or false positive), which can standardize the minimum sequencing depth for different low-frequency mutation detection. Then, the effect of sequencing depth on the fidelity of NA12878 with 3% mutation frequency and circulating tumor DNA (ctDNA of 1%, 3% and 5%) showed that the increase of sequencing depth reduced the fluctuation range of detected-VAFs around the expected VAFs, that is, the fidelity was improved. Finally, based on our experiment result, the consistency of single-nucleotide variants (SNVs) between paired FF and FFPE samples of mice increased with increasing depth, suggesting that increasing depth can improve the precision and sensitivity of low-frequency mutations.

**Highlights:** The normalized relationship between sequencing depth and the probability of true positive mutation (sensitivity) is established based on binomial distribution.

The probability of true positive increases and the probability of false positive decreases when the number of positive mutated reads increases (threshold) in an equal proportion with depth.

Detected-VAFs fluctuates regularly around expected-VAFs. The amplitude of detected-VAFs fluctuation decreases with sequencing depth and the “fidelity” increases.

The increase of “fidelity” leads to a higher degree of differentiation between true and false positive mutations, which ultimately increases the true positive probability and decreases the false positive probability.

## 1. Introduction

As next-generation sequencing (NGS) combined with enrichment technique is becoming an effective method, it is better to accurately detect numerous mutations from high-throughput data in clinical samples[1,2]. Due to the limitations of sample type, sequencing depth and other conditions, how to accurately identify mutations with variant allele frequencies lower than 5% in samples is still a great challenge at present[3]. In addition, the occurrence of most tumors is usually related to somatic mutations (especially single-nucleotide variants, SNVs), and the frequency of these mutations is usually low due to the contamination of normal cells or the heterogeneity of tumors[4]. Therefore, accurate analysis and detection of low-frequency somatic mutations is an urgent problem to be solved. Among them, sequencing depth is a very important factor to achieve accurate detection of low-frequency mutations.

Recently, emerging genetic testing methods, such as liquid biopsy[5] and noninvasive prenatal testing[6], are closely related to the detection of low-frequency somatic mutations and become hot research areas of precision therapy. The two biomarkers in the liquid biopsy are circulating tumor cells (CTCs) and circulating cell-free tumor DNA (ctDNA), which play important roles in the dynamic monitoring of tumor development process[7]. While NGS has limitations in detecting mutation frequencies below 1-2% of ctDNA, the use of targeted NGS technology and gene panels increases the sensitivity by increasing sequencing throughput and depth of the target region[8,9]. Furthermore, since vast clinical material was archived as formalin-fixed paraffin-embedded (FFPE) tissue samples[10], many retrospective studies need to use it for further analysis. However, DNA extracted from FFPE tissues has several types of DNA lesions, such as formaldehyde-induced cross-linking and cytosine base deamination[11,12]. The low-frequency sequencing artifacts caused by these lesions are difficult to distinguish from true mutations[13]. To improve the reliability of FFPE tissue samples as clinical samples, as well as the detection consistency between FFPE and FF, it is necessary to achieve high sensitivity and eliminate the interference of false positive mutations as much as possible[14,15].

It is very important to set reasonable sequencing depth and threshold for detection of true positive mutations with low-frequency. The higher the number of reads compared, the higher the confidence of the mutation, which means that sequencing depth directly affects the recurrence of the mutation detection[16]. And the number of false-positive mutations are largely determined by sequencing errors[17]. The inherent error rate for traditional NGS ranges from 0.1% to 1%, while that of MGISEQ-2000 sequencing platforms was about 0.2%. The insufficient depth and high threshold will reduce the true-positive mutation probability and increase the false negative probability. Undoubtedly, if the threshold of mutated reads is unchanged and only the sequencing depth is increased, the probability of true-positive mutations will definitely increase, but at the same time, the probability of false-positive mutations introduced by sequencing error will also increase accordingly. Therefore, when detecting low-frequency mutations, a reasonable detection threshold must be set to maximize the true positive mutation probability and minimize the probability of false positive and false negative. In conclusion, standardization of sequencing depth and detection threshold is the key to accurate detection of low-frequency mutations[18].

## 2. Material and Methods

### 2.1. True positive probability simulation and sequencing depth standardization based on the binomial distribution

The probability of true positive, false negative and false positive at different sequencing depths and thresholds can be calculated by binomial distribution when the mutation frequency is given[16]. For example, with the binomial distribution, the true positive mutation frequency of 3%, and the sequencing depth of 2500X, the probability of detecting 49 or fewer mutated reads is 0.1%. So, the probability of detecting 50 or more mutated reads is 99.99% (1–0.1%), that is, the probability that true positive mutations of 3% VAF are detected is 99.99% and missed is 0.1%. It can be realized by a simple calculation formula in Excel: BINOM.DIST (49, 2500,3%, true) and 1-BINOM.DIST (49, 2500,3%, true).

Here, we assumed that the true positive frequency was 3% and the false positive frequency was 1%, and set two types of thresholds: fixed number of mutated reads (mutated reads of 100X, 200X, 300X, etc., were 3) and variable mutated reads—the fixed proportion of mutated reads (mutated reads of 100X, 200X and 300X, etc., were 2, 4 and 6 respectively). Finally, the probability of true positive and false positive was obtained (**Table S1** for detail) and the distribution model of detected-VAFs was simulated by BINOM.INV (inverse binomial distribution) (see **Table S2**).

### 2.2. Sample preparation

The NA12878 standard of 3% expected VAF was mixture of the NA12878 and NA24385. The expected VAF of circulating tumor DNA standards in lung cancer were 1%, 3% and 5%, respectively (Gene-Well). we prepared the paired fresh-frozen (FF) and formalin-fixed paraffin-embedded (FFPE) samples from the heart, liver, spleen and kidney of 3 C57BL/6 mice. DNeasy Blood & Tissue Kit and QIAamp DNA FFPE Tissue Kit (Qiagen) were used for Genomic DNA extraction of paired FF and FFPE samples respectively.

### 2.3. Library preparation, hybridization capture and sequencing

The ctDNA of about 180bp was directly used for library preparation. The genomic DNA was sheared (~300bp) by Covaris™ Focused-ultrasonicator (Thermo Fisher Scientific™). 50-100 ng sheared DNA was quantified by Qubit™ 3 Fluorometer (Thermo Fisher Scientific™) and was input for library preparation with MGIEasy universal DNA library kit (MGI). Agilent 2100 Bioanalyzer (Agilent Technologies™) and MGIEasy DNA Clean Beads were respectively used for DNA quality control and purification. MGIEasy Exome Capture Hybridization and Wash Kit were used in hybridization capture of the 3% NA12878 standard, while the NanOnco panel was used in hybridization capture of ctDNA standard and the mice tissues. All captured DNA products were circularized to ssCirDNA, made into the DNA Nanoballs (DNBs) and sequenced with PE100 on the MGISEQ-2000 (MGI) platform.

### 2.4. Sequencing data analysis

Reads which have more than 5 “N” bases, over 50% poly-A and low-quality bases were filtered out by SOAPnuke. The remaining reads were aligned to the human reference genome hg19 or mm10 by BWA-MEM and mapped reads with MAPQ ≥ 10 were used for subsequent analysis. After alignment, single-nucleotide polymorphisms (SNPs) were called by VarScan[19].

NA12878: The distribution of detected VAFs in NA12878 standard was analyzed under 500X sequencing depth and the threshold was set to 0.5%. ctDNA: These ctDNA standards (1%, 3% and 5%) were made in triplicate and sequenced in 3000X depth each. The generated data of ctDNA were analyzed in three sequencing depths (1000X, 2000X and 3000X). To reduce the impact of random downsampling, each depth was randomly downsampled to 20 times. We set two detection thresholds of 0.5% and 0.7% for 1% expected VAF in ctDNA and thresholds for 3% and 5% expected VAF were increased proportionately (i.e., 1%, 3% and 5% expected VAFs correspond to two thresholds respectively, a: 0.5%, 1.5%, 2.5% and b: 0.7%, 2.1%, 3.5%). Then the fluctuation of detected mutation frequency and the sensitivity of ctDNA at different sequencing depths were analyzed. Mice: In addition, we evaluated the SNVs concordance of paired FF and FFPE tissues (heart, liver, spleen and kidney) in mice at the sequencing depth of 500X and 1000X (thresholds: 0.5%). The relationship between SNVs concordance and different sequencing depths was further analyzed.

The data reported in this study are alsoavailable in the CNGB Nucleotide Sequence Archive (CNSA: https://db.cngb.org/cnsa; accession number CNP0002455).

## 3. Results

### 3.1. Binomial distribution: calculation of true/false positive probability and suggestions for sequencing depth standardization

The true and false positive mutation frequencies were assumed to be 3% and 1%, respectively. The specific content of the analysis process is shown in **Method 2.1** and **Table S1-S2**. Threshold of variable mutated reads (2%): the true positive probability still increased with depth, while false positive probability decreases with depth (**Fig. 1A**). Threshold of a fixed number of mutated reads (3 mutated reads): Undoubtedly, the probability of both true and false positive mutations increased with increasing depth (**Fig. 1B**). The puzzle is, why does the probability of true positive still increase and the probability of false positive decrease when the number of positive mutated reads increases in an equal proportion with depth. We simulated and analyzed the distribution of detected-VAFs at different depths through BINOM.INV (inverse binomial distribution), and it is found that the higher the depth, the closer detected-VAFs to the theoretical VAFs (3% and 1%), that is, the higher the fidelity of detected-VAFs (**Fig. 1C**). This phenomenon is the embodiment of the law of large numbers. In **Fig. 1C**, when the threshold is 2%, the increase of fidelity leads to a higher degree of discrimination between true and false positive mutations, as many true positive mutations as possible are detected (true positive probability increase), while false positive less than the detection threshold are filtered (false positive probability decrease).

**Fig. 1.**
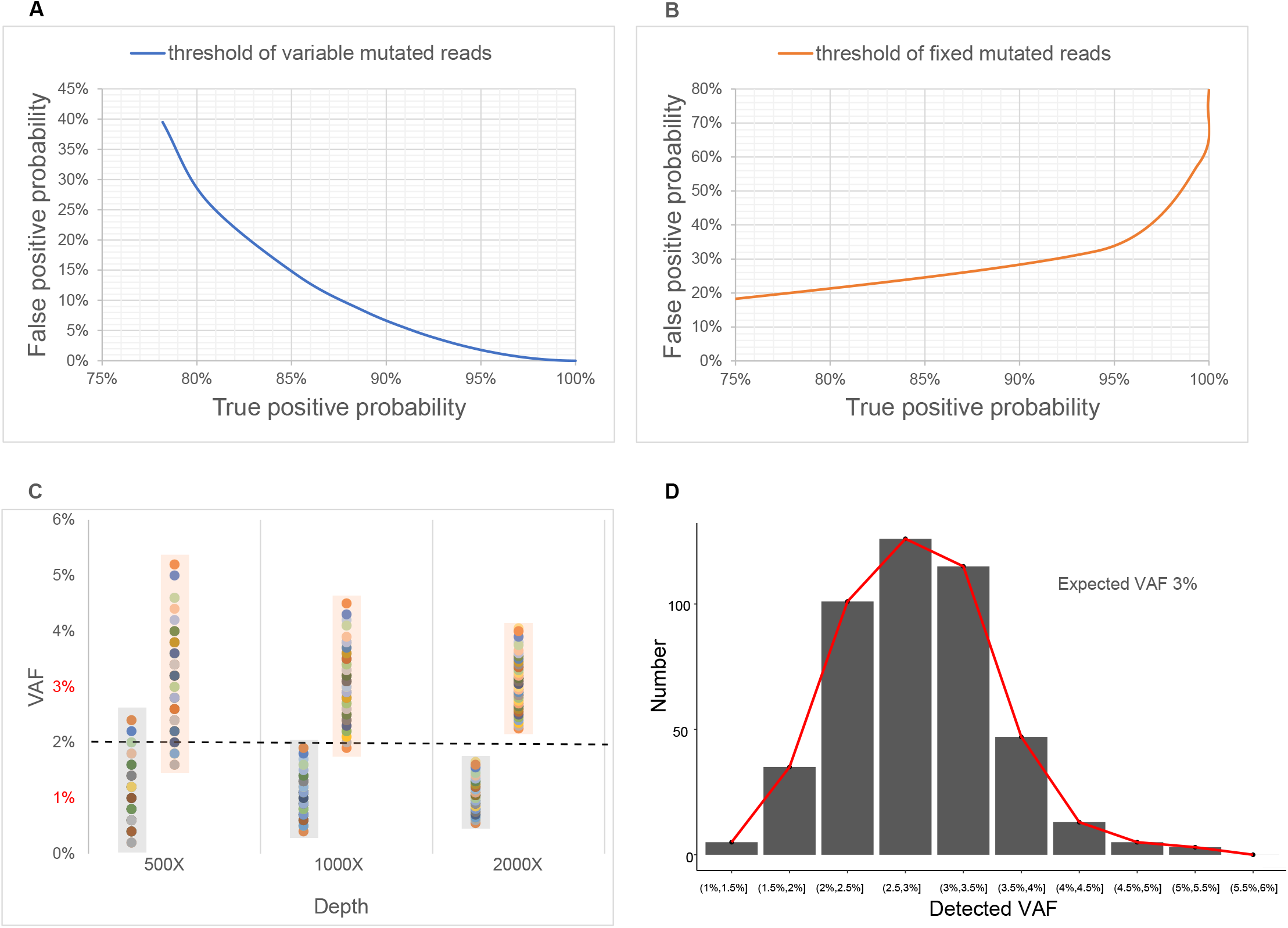
Calculation of true/false positive probability by binomial distribution and the distribution of detected-VAFs. The expected-VAFs of true and false positive mutation was assumed to be 3% and 1%, respectively. (A) The true/false positive probability varies with sequencing depth when the threshold is 2%. (B) The true/false positive probability varies with depth when the threshold is 3 positive mutated reads. (C) Distribution of detected-VAFs at different depths through inverse binomial distribution when the threshold is 2%. (D) The histogram and curve distribution of detected-VAFs in NA12878 with expected-VAFs of 3%. Detected-VAFs were divided into certain intervals and the number of mutations in each interval was counted to obtain the normal distribution.

### 3.2. Mutation detection of NA12878 standard

For NA12878 with expected-VAFs of 3%, to more intuitively display the varying regularity of detected-VAFs at a certain sequencing depth, we divided several frequency intervals, counted the number of mutations in each interval, and obtained the normal distribution of detected-VAFs as shown in **Fig. 1D**. The mutation detection result clearly reflects the fluctuation of the detected-VAFs around expected-VAFs, which ranges from 1% to 5%. Both library preparation and sequencing processes affected the detection of mutation frequency and caused detected-VAFs to fluctuate around expected-VAFs. The regular fluctuation of mutation frequency within a certain range in actual detection is consistent with the result of simulated data in **Fig. 1C**.

### 3.3. Distribution of detected-VAFs in ctDNA standards at different depths

The mutation site information and summary of ctDNA samples were shown in **Table S3 and S4**. We set two thresholds for ctDNA with expected-VAFs of 1%, 3% and 5% respectively (a: 0.5%, 1.5%, 2.5% and b: 0.7%, 2.1%, 3.5%) to analyze the relationship between detected-VAFs and expected-VAFs at different depths (**Method 2.4**). With the increase of sequencing depth, the floating range of the detected-VAFs decreased significantly and became closer to corresponding expected-VAFs (**Fig. 2A**). The result persisted even when the threshold was increased (**Fig. 2B**). We called that the detected-VAFs got high “fidelity” at high sequencing depth. The order of fluctuation amplitude of different sequencing depths was: 3000X<2000X<1000X. In addition, according to the above simulation results of the binomial distribution, the improvement of fidelity is bound to increase the sensitivity of mutation detection. It was evident from **Table 1** that the sensitivity increased with sequencing depth. When the sequencing depth was 2000X, the sensitivity of expected-VAFs of 3% or 5% was 100%, while the sensitivity of expected-VAFs of 1% was only 87% at 3000X (threshold: a, **Table 1**).

**Fig. 2.**
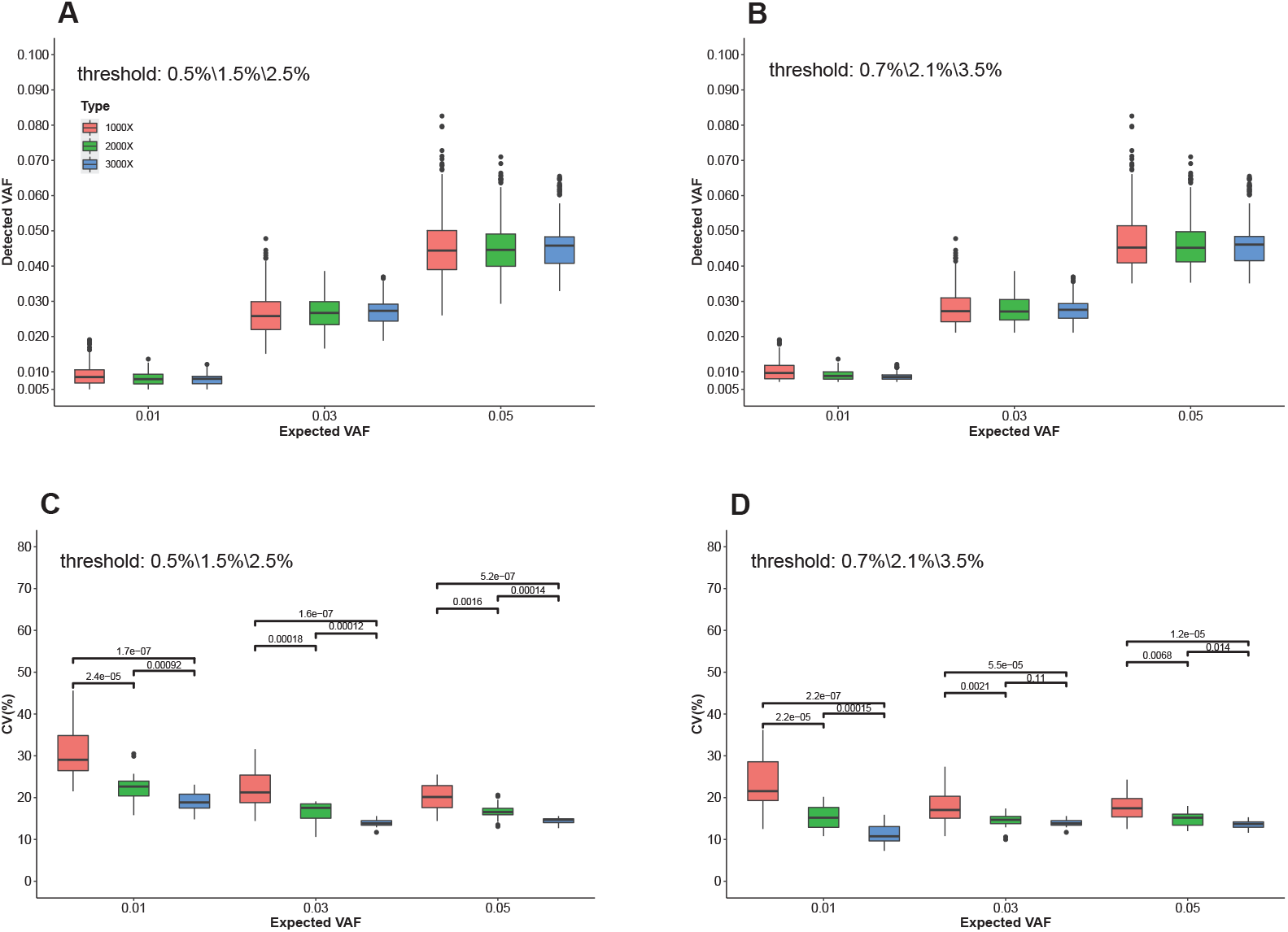
Box plot of detected VAFs at different depths in the ctDNA samples. (A) The red, green, and blue boxplot represent three sequencing depths: 1000X, 2000X, and 3000X, respectively. Distribution of detected VAFs at different depths when the threshold was set at 0.5%, 1.5% and 2.5% for the expected VAFs of 1%, 3% and 5%, respectively. (B) Distribution of detected VAFs at different depths when the threshold was set at 0.7%, 2.1% and 3.5% for the expected VAFs of 1%, 3% and 5%, respectively. The coefficient of variations (CVs) measured the fluctuation range of detected VAFs at different depths when the thresholds were 0.5%, 1.5% and 2.5% (C); or the thresholds were 0.7%, 2.1% and 3.5% (D).

**Table 1:**
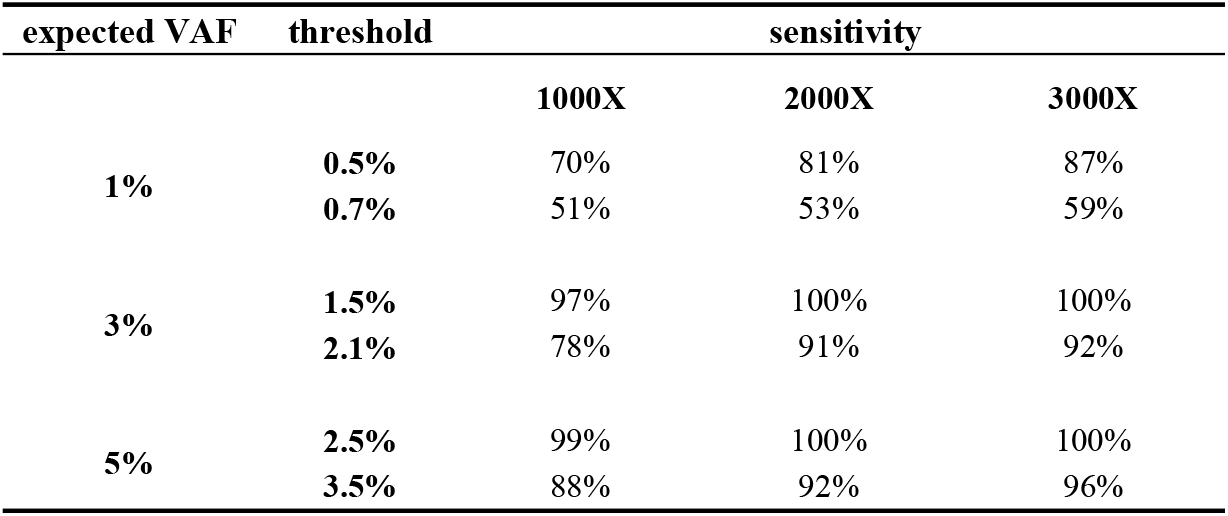
Detection sensitivity at different sequencing depths (ctDNA with expected-VAFs of 1%, 3% and 5%)

On the other hand, the fluctuation range of detected-VAFs was measured by the coefficient of variations (CVs) at different depths. It can be seen that the CVs decreased with depth (**Fig. 2C and 2D**). The Wilcoxon signed-rank test confirmed that there was statistically significant difference between the CVs and different sequencing depths, and there were significant negative correlations between the two (p-value is less than or equal to 0.05). Finally, to verify the accuracy of the mutation detection results, the mutation frequencies detected by ddPCR were compared (**Fig. S5)**.

### 3.4. Single-nucleotide Variants (SNVs) concordance between paired FF and FFPE samples in mouse

Unlike the NA12878 and ctDNA standards, the expected-VAFs (true mutation frequencies) of the real samples were unknown, so it cannot directly reflect the relationship between detected-VAFs and expected-VAFs in the real samples. Therefore, the SNVs consistency of paired FF and FFPE tissues in mice at different sequencing depths was used to imply that the fidelity of detected-VAFs and the sensitivity increased with sequencing depth. All SNVs in each tissue were analyzed at 500X and 1000X with the threshold of 0.5% (**Fig. 3, Fig. S1-S2**).

**Fig. 3.**
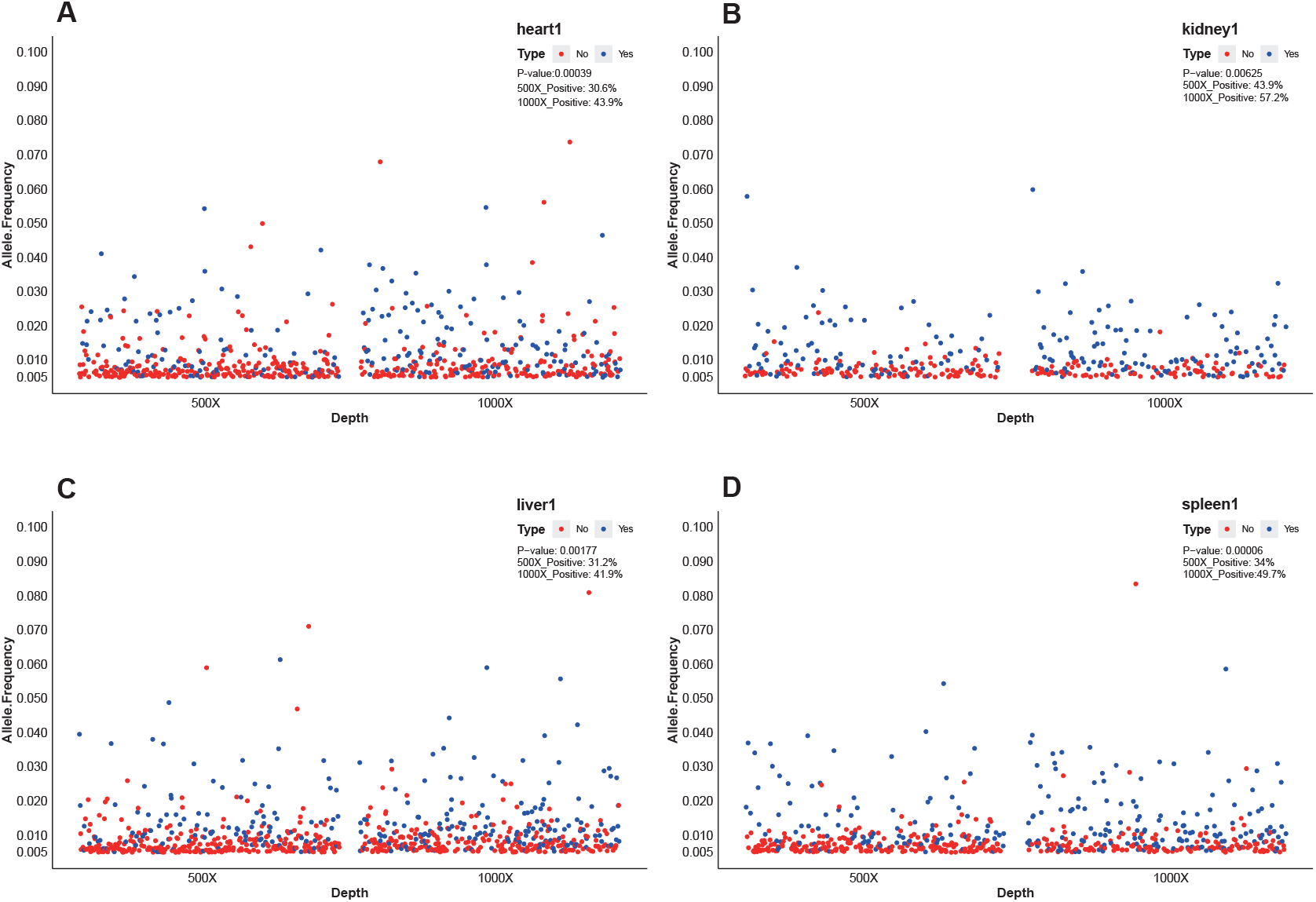
Single-nucleotide Variants (SNVs) concordance between paired FF and FFPE samples at different depths in mouse1. (A) The mutation frequency distribution of all SNVs in the FFPE sample of heart1 was analyzed at 500X and 1000X. The red and blue dots in each FFPE tissue sample represented SNVs that were inconsistent with FF and consistent with FF respectively. The mutation consistency of FF and FFPE tissue samples from the heart1 was significantly different at 500X and 1000X, as was the case with other tissue samples (p-value less than 0.05 is statistically significant). 500X_positive: the consistency rate (positive rate) of each pair of FF and FFPE tissues at 500X. 1000X_positive: the consistency rate (positive rate) of each pair of FF and FFPE tissues at 1000X. Mutation frequency distribution and concordance of all SNVs FFPE samples of kidney1(B), liver1(C) and spleen1(D) were analyzed at 500X and 1000X.

Here we only show SNVs with variant allele frequencies less than 10%, because they account for more than 90%. The concordance was calculated as a percentage of FFPE mutations that were also identified in the FF samples. The consistency rate (positive rate) of each pair of FF and FFPE tissues at 1000X depth increased by more than 10% compared to 500X depth (**Fig. 3**). The Chi-square test showed that the mutation consistency of each pair of FF and FFPE tissue samples was significantly different at 500X and 1000X. Theoretically, variants concordance should be high because most of the mutations in the FFPE sample should also remain in the FF sample, but the concordance in **Fig. 3** was poor. In addition, the VAFs of discordant mutations mainly distributed in the range of 0.5%~1% and C: G>T: A base substitutions accounted for a high proportion (predominant sequence artefacts in FFPE samples[20,21]), Which suggested the existence of false-positive mutations in the result. With the increase of sequencing depth, the fidelity and true positivity probability increased, and the discrimination of true and false positive mutations was improved as well as the sensitivity, thus increasing the mutation consistency of FF and FFPE.

## 4. Discussion

Sequencing depths and different low-frequency mutations have great impacts on the accuracy, and increasing sequencing depth is a widely used method to improve the accuracy of mutation detection[22]. Targeted next-generation sequencing can achieve higher depth and coverage for a specific area to make the detection of low-frequency mutations more accurate and is widely used in the screening and analysis of somatic mutations in some clinical cancer samples[23]. However, most sequencing data is analyzed for mutations of ≥5% VAFs and it is also a great challenge to achieve accurate and sensitive detection of mutations with VAF below 5% [3].

Additionally, setting a reasonable detection threshold is also crucial. Two parameters, the minimum sequencing depth and detection threshold, are required to evaluate accurate detection of low-frequency mutations. Here, we studied the influence of sequencing depth on the fidelity and sensitivity of 1%-5% low-frequency mutation detection and suggested standardization of sequencing depth calculation.

Here, we found that when the number of positive mutated reads increased in an equal proportion with depth, the true positive probability still increased with depth, while false positive probability decreased with increasing depth. Through binomial distribution simulation and experimental test, it was found that the “fidelity” of detected-VAF was the cause of this phenomenon. For NA12878 with expected-VAFs of 3%, the normal distribution of the detected-VAFs showed more visually that the detected-VAFs were floating regularly around the expected VAFs. In addition, detected-VAF of ctDNA (lung cancer) positive standard with defined mutation sites and mutation frequency was more concentrated with depth, which confirmed our conclusion that low frequency mutation detection at higher depth was more real and accurate. And this finding was also applicable to real samples because of the significant relationship between the sequencing depth and the mutation consistency of FF and FFPE. Therefore, we emphasize the importance of determining sequencing depth in low-frequency mutations to obtain a credible, reproducible assay. Achieving accurate detection of low-frequency mutations and standardization of sequencing depth is a challenge with profound clinical implications[16,24].

Nevertheless, some sequencing errors cannot be ignored in the analysis process. The sequencing errors may come from many stages during DNA processing, library preparation and or special samples such as FFPE. Therefore, a reasonable detection threshold should also be set based on sequencing errors to minimize false-positive mutations. Our experimental results are based on MGISEQ-2000, and the sequencing error is about 0.2%. UMI (Unique Molecular Identifiers) are used as an effective error correction method for PCR errors[25].

In conclusion, with the increase of sequencing depth (positive mutated reads increased in an equal proportion with depth), the fidelity, true positivity probability and sensitivity were improved. Minimum sequencing depth can also be calculated by binom. dist based on the true positive mutation probability to be achieved, making the required depth more standard.

## 6. Acknowledgments

We thank all members of the Center for Application R&D from MGI for helpful comments and Henan Key Laboratory for Pharmacology of liver diseases for assistance in mice sample collection.

## 7. Author contributions statement

ZL, WJQ and SJF conceived the idea. WJQ analyzed the data with the assistance of ZL and SJF. ZL completed the experiment with the assistance of SJF and MZL. ZL wrote and revised the manuscript with the guidance of JX, XZ and ZLL. CYG and YQY provided helpful comments on this study. All authors reviewed and approved the final manuscript.

## Supplementary figure legends

**Fig.S1**: Single-nucleotide Variants (SNVs) concordance between paired FF and FFPE samples at different depths in mouse2.

**Fig.S2**: Single-nucleotide Variants (SNVs) concordance between paired FF and FFPE samples at different depths in mouse3.

## Supplementary table legends

**Table S1:** Calculation of true/false positive probability by the binomial distribution.

**Table S2:** The distribution model of detected-VAFs at different sequencing depths by the inverse binomial distribution.

**Table S3:** Standard mutation site information of ctDNA.

**Table S4:** Summary of ctDNA samples.

**Table S5:** The mutation frequencies detected by ddPCR.

