## Supplementary figures and images for "Influence of sequencing depth on the fidelity and sensitivity of 1%-5% low-frequency mutation detection and recommendation for standardization of sequencing depth"

### Fig. S1

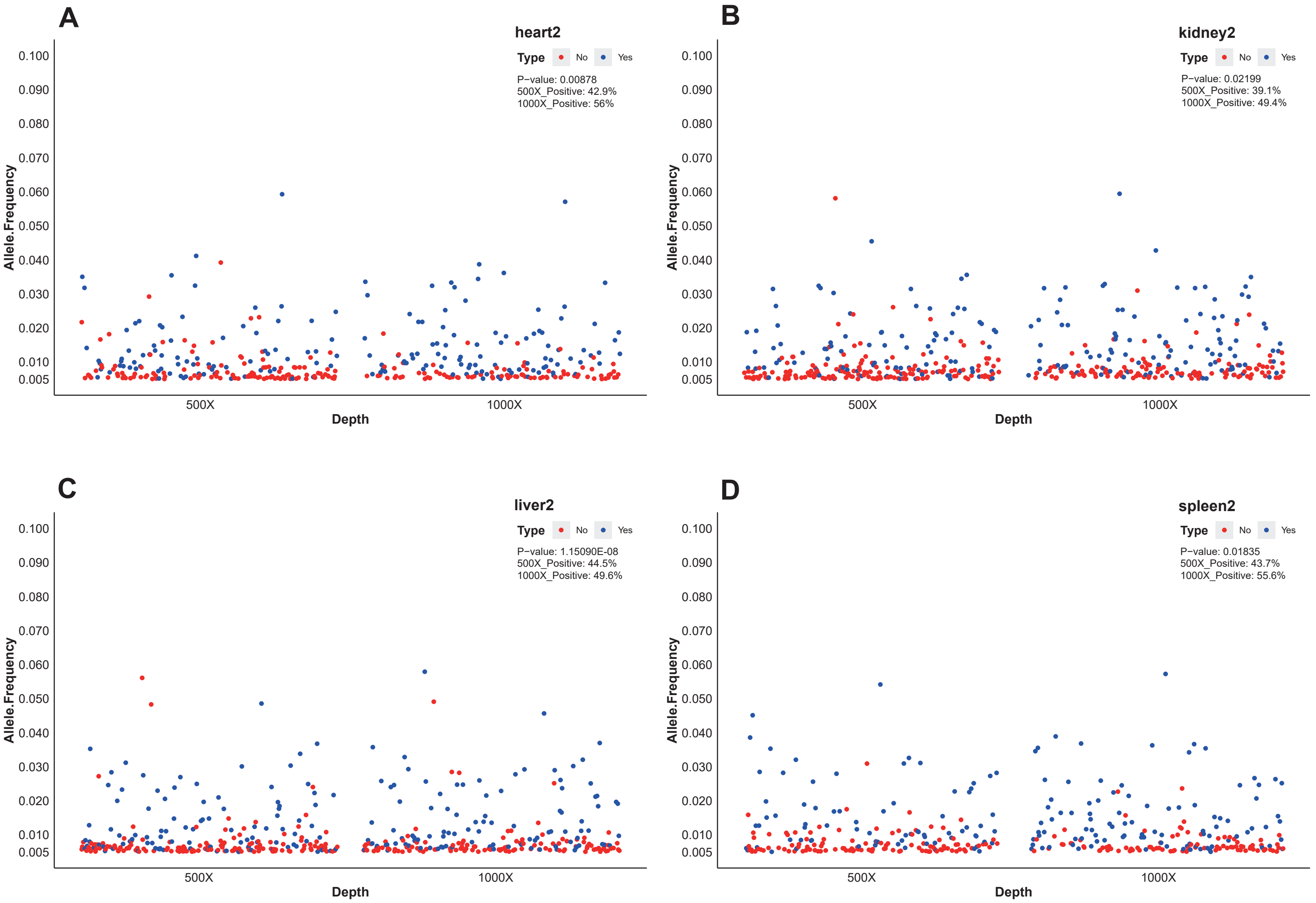

### Fig. S2

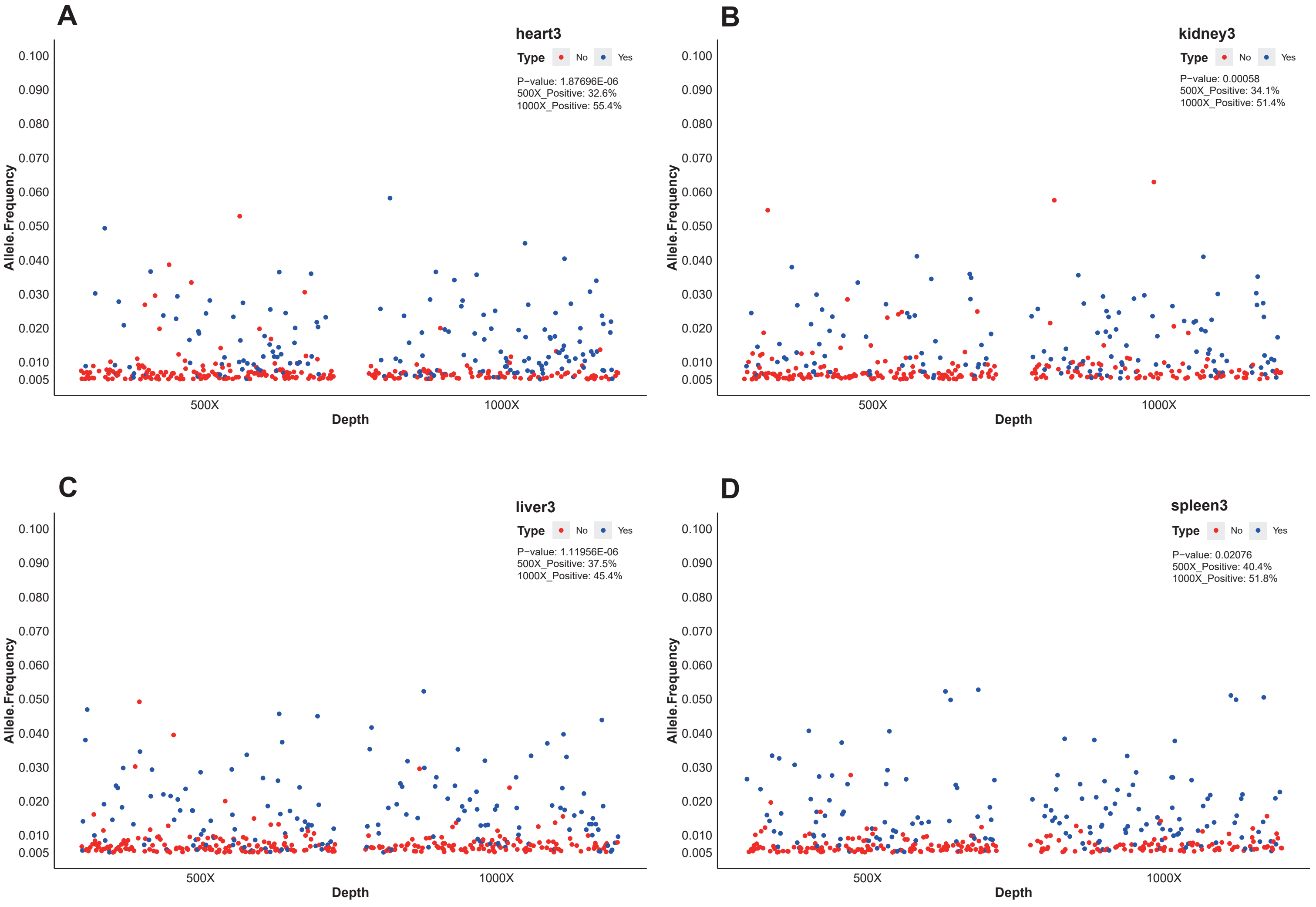
